# Specific hippocampal interneurons shape consolidation of recognition memory

**DOI:** 10.1101/842021

**Authors:** Jose F. Oliveira da Cruz, Arnau Busquets-Garcia, Zhe Zhao, Marjorie Varilh, Gianluca Lavanco, Luigi Bellocchio, Laurie Robin, Astrid Cannich, Francisca Julio-Kalajzić, Filippo Drago, Giovanni Marsicano, Edgar Soria-Gomez

## Abstract

A complex array of different inhibitory interneurons tightly controls hippocampal activity, but how such diversity specifically impacts on memory processes is scantly known. We found that a small subclass of type-1 cannabinoid receptor (CB_1_)-expressing hippocampal interneurons determines episodic-like memory consolidation by linking dopamine D_1_ receptor signaling to GABAergic transmission.

Mice lacking CB_1_ in D_1_-positive cells (D_1_-*CB_1_*-KO) displayed impaired long-term, but not short-term, object recognition memory. Re-expression of CB_1_ in hippocampal, but not striatal, D_1_-positive cells rescued this memory impairment. Learning induced a facilitation of *in vivo* hippocampal long-term potentiation (LTP), which was abolished in mutant mice. Chemogenetic and pharmacological experiments revealed that both CB_1_-mediated memory and associated LTP facilitation involves the local control of GABAergic inhibition in a D_1_-dependent manner.

This study reveals that CB_1_-/D_1_-expressing interneurons shape hippocampal circuits to sustain recognition memory, thereby identifying a mechanism linking the diversity of hippocampal interneurons to specific behavioral and cognitive outcomes.

## INTRODUCTION

The formation of episodic memory is a multistep brain process that requires the coordinated and sequential activity of different brain structures of the medial temporal lobe (Squire et al., 2007). In particular, the hippocampus plays a key role in memory consolidation allowing the long-term storage of recently acquired events. Hippocampal circuits are regulated by a large variety of local inhibitory interneurons, whose identities and functions are under intense scrutiny (Harris et al., 2018, Pelkey et al., 2017, Parra et al., 1998). These interneurons are controlled by complex neuromodulatory systems ensuring their coordinated functions to shape behavioral responses (Klausberger and Somogyi, 2008).

The endocannabinoid system is a major brain modulatory signaling hub mainly formed by type-1 cannabinoid (CB_1_) receptors, their endogenous ligands (endocannabinoids) and the enzymatic machinery for the synthesis and degradation of endocannabinoids. In the hippocampus, CB_1_ receptors are present in principal neurons and astroglial cells (Busquets-Garcia et al., 2015, Oliveira da Cruz et al., 2016). However, the far largest expression of CB_1_ receptors resides in a subgroup of GABAergic interneurons (Marsicano and Kuner, 2008, Katona and Freund, 2012), where they modulate local inhibition of hippocampal circuits. Particularly, the largest amount of CB_1_ receptors is expressed in cholecystokinin (CCK)-positive cells, which constitute a heterogeneous class of GABAergic interneurons characterized by asynchronous neurotransmitter release (Harris et al., 2018, Katona et al., 1999).

Hippocampal CB_1_ receptors are involved in numerous cognitive functions. For instance, CB_1_ receptors control episodic-like memory processes and hippocampal synaptic plasticity *in vivo* and *ex vivo* (Monory et al., 2015, Robin et al., 2018, Hebert-Chatelain et al., 2016, Puighermanal et al., 2009). However, the specific locations where these functions are exerted are far from being fully understood and their investigation is required to characterize the mechanisms of hippocampal-dependent memory.

Activity-dependent long-term changes in hippocampal synaptic transmission are considered cellular correlates of learning and memory consolidation (Nicoll, 2017, Whitlock et al., 2006), which involves local D_1_ receptor signaling (Lisman et al., 2011, Yamasaki and Takeuchi, 2017). For instance, the exposure to hippocampal-dependent behavioral tasks induces changes in long-term potentiation (LTP) of synaptic transmission that require the activation of D_1_-like receptors (Frey et al., 1990, Granado et al., 2008, Li et al., 2003, Lemon and Manahan-Vaughan, 2006). Recently, it has been shown that a subclass of hippocampal CCK-positive interneurons, which express CB_1_ receptors, contains dopamine D_1_ receptors, thus potentially representing a novel subpopulation of CB_1_ expressing hippocampal interneurons (Puighermanal et al., 2017, Gangarossa et al., 2012). However, the potential interactions between D_1_ and CB_1_ receptors in regulating learning-induced plasticity, activity of hippocampal inhibitory circuits and memory processes remain unexplored.

Using a combination of conditional and virally-induced mutagenesis, behavioral analyses, chemogenetics, pharmacology and *in vivo* electrophysiological approaches, here we assessed the role of CB_1_ receptors expressed in D_1_-positive cells in the regulation of episodic-like object recognition memory. We found that conditional deletion of the *CB_1_* gene in D_1_-positive cells impairs long-, but not short-term recognition memory. Strikingly, this function of CB_1_ receptors is triggered by endogenous activation of D_1_ receptors in a small subpopulation of hippocampal interneurons co-expressing the two receptors. Thus, CB_1_ receptor signaling provides a functional link between hippocampal dopaminergic and GABAergic control of synaptic plasticity and memory consolidation.

## RESULTS

### CB_1_ RECEPTORS IN HIPPOCAMPAL D_1_-POSITIVE NEURONS ARE NECESSARY FOR THE CONSOLIDATION OF OBJECT RECOGNITION MEMORY

To evaluate the role of CB_1_ receptors in D_1_-positive neurons, we tested mutant mice bearing a deletion of the *CB_1_* receptor gene in cells expressing D_1_ receptors (D_1_-*CB_1_*-KO mice) (Monory et al., 2007) in the L-maze version of the novel object recognition memory task (NOR, **Figure 1A**) (Puighermanal et al., 2009, Busquets-Garcia et al., 2011, Robin et al., 2018). While D_1_-*CB_1_*-KO mice displayed no phenotype in the short-term version of the task (3h after training, **Figure 1B**), they showed a strong impairment in long-term (24h) memory as compared to their wild-type (WT) littermates (**Figure 1C**), with no changes in total exploration time (**Figure S1A-S1D**).

**FIGURE 1.**
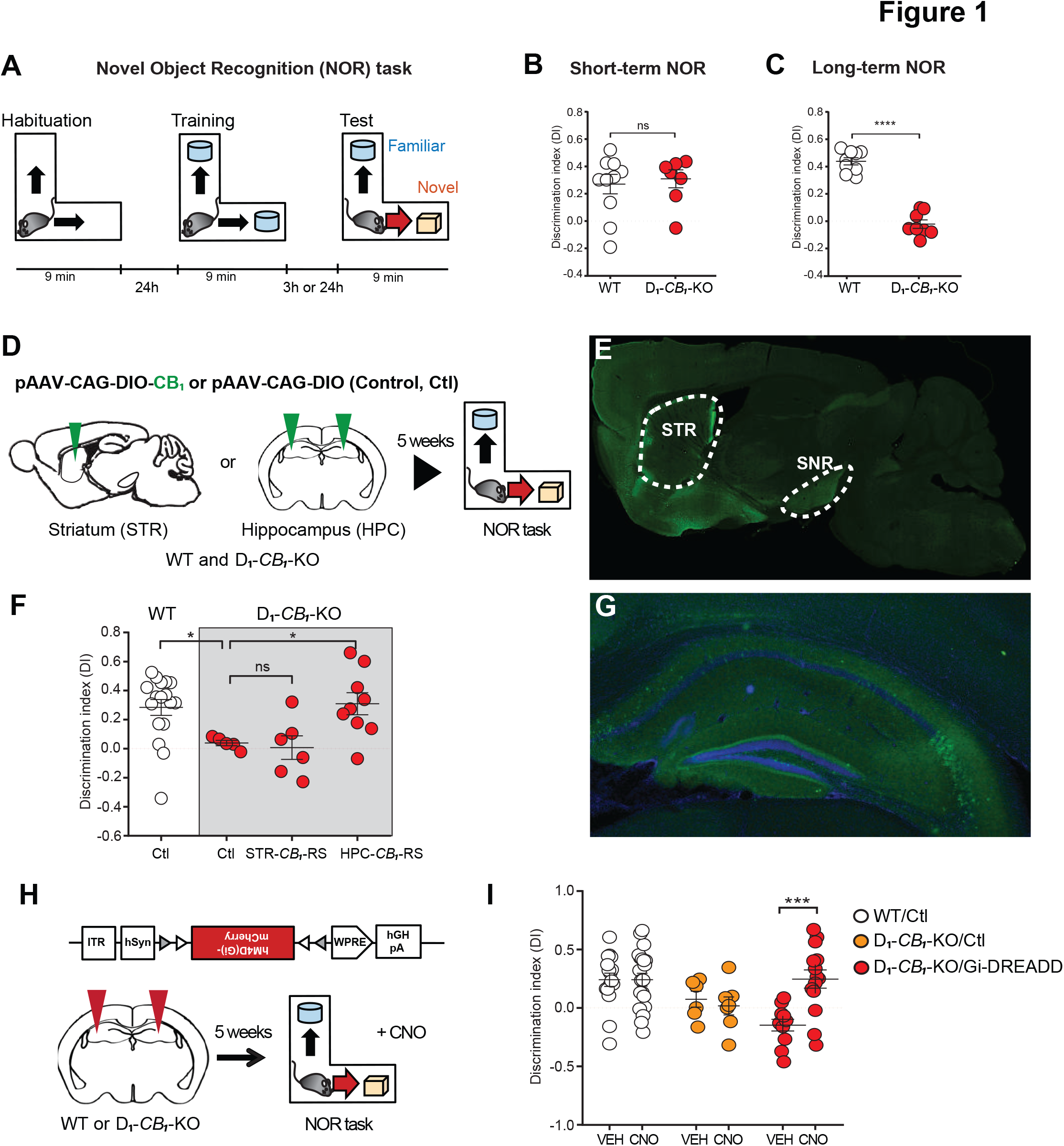
HIPPOCAMPAL CB_1_ IN D_1_-POSITIVE CELLS ARE NECESSARY FOR LATE, BUT NOT EARLY, CONSOLIDATION OF OBJECT RECOGNITION MEMORY. (A) Schematic representation of the NOR Memory task. (B) Memory performance of D_1_-*CB_1_*-WT mice (n = 10) and D_1_-*CB_1_*-KO littermates (n = 7) in the short-term version of the NOR task (3 hours delay between the Training and the Testing phase). (C) Memory performance of D_1_-*CB_1_*-WT mice (n = 9) and D_1_-*CB_1_*-KO littermates (n = 8) in the long-term version of the NOR task (24 hours delay between the Training and the Testing phase). (D) Schematic representation of the experimental design. 5 weeks after the local, CRE-dependent (i.e. D_1_-expressing cells) viral re-expression of the *CB_1_* gene in the striatum (STR) or the hippocampus (HPC) of D_1_-*CB_1_*-WT mice and D_1_-*CB_1_*-KO littermates, mice are tested in the NOR task. (E) NOR memory performance of mice intra-hippocampally injected with Control [n(D_1_-*CB_1_*-WT)=17 and n(D_1_-*CB_1_*-KO)=5], STR*-CB_1_*-RS [n(D_1_-*CB_1_*-KO)=6] or HPC-*CB_1_*-RS [n(D_1_-*CB_1_*-KO)=9]. (F and G) Representative images of slice of CRE-expressing D_1_-*CB_1_*-KO mice injected with CB_1_-myc in the STR or HPC using the same procedure as described in (D) (see methods). (F) Immunofluorescence of neurons expressing CB_1_-myc from mice injected with the virus in the striatum. (G) Immunofluorescence of neurons expressing CB_1_-myc from mice injected with the virus in the hippocampus. (H) Schematic representation of the experimental design. 5 weeks after the local, CRE-dependent (i.e. D_1_-expressing cells) viral expression of the Gi-DREADDs or mCherry in hippocampus of D_1_-*CB_1_*-WT mice and D_1_-*CB_1_*-KO littermates, mice are tested in the NOR task. Clozapine N -oxide (CNO) injections take place immediately after the Training phase of the NOR task. (I) NOR memory performance of D_1_-*CB_1_*-WT mice intra-hippocampally injected with hM4D(Gi) virus or mCherry [n(VEH)=16 and n(CNO)=21], D_1_-*CB_1_*-KO mice intra-hippocampally injected with mCherry [n(VEH)=6 and n(CNO)=7] and D_1_-*CB_1_*-KO mice intra-hippocampally injected with hM4D(Gi) [n(VEH)=11 and n(CNO)=14]. Data, mean ± SEM. *p < 0.05, ***p < 0.001, ****p < 0.0001, ns=not significant. See also **Figure S1** and **Table S1**.

The majority of CB_1_ receptors in D_1_-positive neurons have been previously characterized in striatonigral circuits (Monory et al., 2007). Considering that these circuits have been implicated in object recognition memory (Darvas and Palmiter, 2009), we first tested the role of striatal CB_1_ receptors in our NOR task. We infused an adeno-associated virus carrying a Cre-dependent expression of CB_1_ (pAAV-CAG-DIO-*CB_1_*) into the striatum of D_1_-*CB_1_*-KO mice to obtain a re-expression of CB_1_ in D_1_-positive cells of this brain region (STR-*CB_1_*-RS mice, **Figure 1D,E**). This manipulation induced a reliable re-expression of the CB_1_ receptor in D1-positive striatonigral projecting neurons as revealed by the parallel immunodetection of a myc-tagged version of the CB_1_ protein (CB_1_-myc see Methods, **Figure 1E**). However, this re-expression was not sufficient to rescue the phenotype of D_1_-*CB_1_*-KO mice in long-term NOR (**Figure 1F**, **S1E, S1F**). This suggests that CB_1_ receptors in striatal D_1_-positive cells do not participate to this type of memory. Anatomical data indicate that a subset of hippocampal neurons contain D_1_ receptors (Gangarossa et al., 2012), likely co-expressing CB_1_ protein (Puighermanal et al., 2017). As the hippocampus is a key region for memory, we used the same strategy as described above, to re-express the *CB_1_* gene in the hippocampus of D_1_-*CB_1_*-KO mice to obtain HPC-*CB_1_*-RS mice (**Figure 1D,G**). This manipulation fully rescued the phenotype of the mutant mice (**Figure 1F,G**; **S1E and S1F**). These data indicate that hippocampal CB_1_ receptors expressed in D_1_-positive cells are required for NOR memory.

We have recently reported that deletion of CB_1_ receptors in hippocampal glial acidic fibrillary protein (GFAP)-positive cells (i.e. mainly astrocytes, GFAP-*CB_1_*-KO mice) impaired the consolidation of NOR memory in the same task (Robin et al., 2018). Indeed, GFAP-*CB_1_*-KO mice were strongly impaired in NOR (**Figure S1G-S1I**), confirming previous results (Robin et al., 2018). However, in contrast to D_1_-*CB_1_*-KO mice, the deletion of CB_1_ in astroglial cells also impaired short-term NOR memory (**Figure S1J-S1L**). This difference indicates that CB_1_ receptors expressed in hippocampal astrocytes or D_1_-positive cells control earlier and later consolidation phases of object recognition, respectively. Thus, specific hippocampal CB_1_ receptor subpopulations subserve distinct roles in the same memory process.

The primary function of CB_1_ receptor activation in neurons is to decrease neurotransmitter release (Castillo et al., 2012, Busquets-Garcia et al., 2017). Accordingly, the deletion of CB_1_ receptors from neurons often results in excessive neurotransmission. Thus, we reasoned that artificial inhibition of hippocampal D_1_-positive neurons during consolidation should be able to rescue the memory impairment of D_1_-*CB_1_*-KO mice. Viral vectors carrying CRE-dependent expression of an inhibitory Designer Receptor Exclusively Activated by Designer Drugs (DIO-hM4DGi, DREADD-Gi) (Robinson and Adelman, 2015) or control mCherry protein were infused into the hippocampi of D_1_-*CB_1_*-KO mice and control WT littermates (**Figure 1H**). Post-training CNO injections (2 mg/kg, intraperitoneal) did not affect the NOR performance of WT mice injected with either DREADD-Gi or mCherry, indicating that the drug or its metabolites had no effect *per se* [(Gomez et al., 2017), **Figure 1I**, **S1M and S1N**]. Similarly, neither vehicle nor CNO injections had any effect in D_1_-*CB_1_*-KO mice infused with a control virus expressing only mCherry (**Figure 1I**, **S1M and S1N**). Conversely, post-acquisition CNO treatment fully rescued the NOR memory impairment of D_1_-*CB_1_*-KO mice expressing DREADD-Gi (**Figure 1I**; **S1M and S1N**). This strongly suggests that excessive activity of D_1_-positive neurons during the consolidation process is responsible for the memory impairment observed in D_1_-*CB_1_*-KO mice.

### CB_1_ RECEPTORS IN HIPPOCAMPAL D_1_-POSITIVE NEURONS CONTROL LEARNING-INDUCED CHANGES OF LTP IN VIVO

Cellular and molecular mechanisms underlying activity-dependent changes in synaptic plasticity are proposed to participate in the formation of long-term memory (Aggleton and Morris, 2018). Previous studies have shown that conditional and global deletion of CB_1_ receptors in neuronal and glial cell populations induces deficits in learning and associated synaptic plasticity (Busquets-Garcia et al., 2017). To address the role of CB_1_ receptors in hippocampal D_1_-positive neurons in the modulation of synaptic plasticity, we recorded *in vivo* evoked field excitatory postsynaptic potentials (fEPSPs) in the hippocampal CA3-CA1 pathway of anesthetized mice. Surprisingly, High Frequency Stimulation (HFS) was able to induce a long-lasting LTP of synaptic fEPSPs in both D_1_-*CB_1_*-KO and WT littermates (**Figure 2A, 2B**), indicating that CB_1_ receptors expressed in hippocampal D_1_-positive neurons are not necessary for the expression of LTP in naïve animals.

**FIGURE 2.**
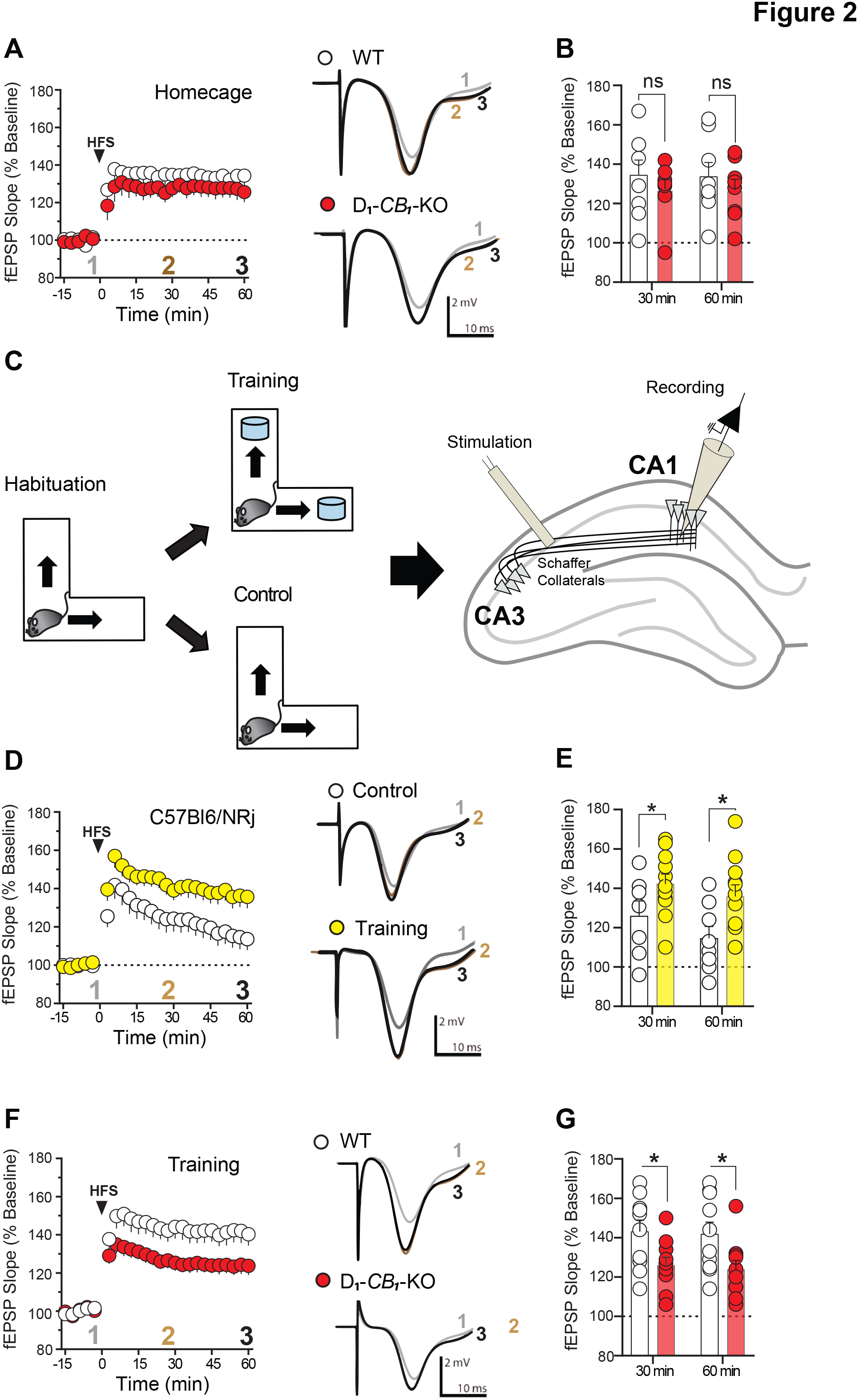
LEARNING-INDUCED FACILITATION OF *IN VIVO* HIPPOCAMPAL LTP REQUIRES CB_1_ RECEPTORS AT D_1_-POSITIVE NEURONS. (A and B) Induction of high-frequency stimulation in the dorsal hippocampal CA3 Shaffer-Collateral pathway induces an *in vivo* LTP in dorsal CA1 *stratum radiatum*. (A) Summary plots of recorded evoked fEPSP in anesthetized D_1_-*CB_1_*-WT (n=8) and D_1_-*CB_1_*-KO (n=8) mice. (B) Bar histograms of normalized fEPSP from (A) representing 30 and 60 minutes after HFS. (C) Schematic representation of the experimental setup. In the first day, naïve C57BL6/NRj are habituated to the NOR L-maze. 24 hours later, animals are exposed to the maze either with two identical objects (i.e. learning, NOR Training) or without objects (Control). Then, right after the behavioral task, mice are anesthetized and place under the *in vivo* electrophysiology procedures (see methods). (D and E) Learning modulates *in vivo* LTP. (D) Summary plots of recorded evoked fEPSP from mice expose to Control (n=8) and NOR Training (n=11) conditions. (E) Bar histograms of normalized of evoked fEPSP from (D) representing 30 and 60 minutes after HFS. (F and G) Learning-induced modulation of *in vivo* LTP is impaired in D_1_-*CB_1_*-KO mice. (F) Summary plots of recorded fEPSP in anesthetized D_1_-*CB_1_*-WT (n=10) and D_1_-*CB_1_*-KO (n=10) mice. (G) Bar histograms of normalized of evoked fEPSP from (F) representing 30 and 60 minutes after HFS. Traces on the right side of the summary plots represent 150 superimposed evoked fEPSP before HFS (1, grey), 30 minutes (2, brown) and 60 minutes (3, black) after HFS. Data, mean ± SEM. *p < 0.05, ns=not significant. See also **Table S1**.

Hippocampal-dependent memory-like processes such as LTP are sensitive to pharmacological and genetic modulation of hippocampal D_1_ receptors, particularly after learning events (Li et al., 2003, Lemon and Manahan-Vaughan, 2006, Takeuchi et al., 2016, Yamasaki and Takeuchi, 2017). Thus, we hypothesized that CB_1_ receptors in D_1_-positive neurons may modulate learning-dependent hippocampal synaptic plasticity. To explore whether acquisition of the NOR task modulates *in vivo* LTP, we exposed C57Bl6/NRj mice to the L-maze containing two identical objects (NOR acquisition) or no object (control condition) and then recorded *in vivo* fEPSP (**Figure 2C**). Interestingly, the HFS of Shaffer collateral/CA1 pathway induced stronger LTP in animals exposed to NOR acquisition than in control mice (**Figure 2D, E**), showing that the training in the NOR task can modulate synaptic plasticity in the hippocampus. Strikingly, D_1_-*CB_1_*-KO mice lacked this learning-induced enhancement of LTP (**Figure 2F, G**). Thus, physiological activation of CB_1_ receptors in hippocampal D_1_-positive neurons is required for a behavioral-dependent facilitation of *in vivo* LTP.

### CB_1_ RECEPTORS IN D_1_-POSITIVE NEURONS MODULATE NOR MEMORY CONSOLIDATION THROUGH A GABAERGIC-DEPENDENT MECHANISM

D_1_ receptors are expressed in different subpopulations of hippocampal cells, including subsets of GABAergic interneurons and pyramidal neurons (Gangarossa et al., 2012). Considering that CB_1_ receptor signaling decreases the activity of both excitatory and inhibitory hippocampal neurons (Busquets-Garcia et al., 2017, Castillo et al., 2012), we asked whether excessive glutamatergic or GABAergic neurotransmission might underlie the phenotype of D_1_-*CB_1_-*KO mice. To address this issue, we injected non-amnesic doses (Puighermanal et al., 2009) of the NMDA receptor blocker MK-801 (0.1 mg/kg, i.p.) or of the GABA_A_ receptor antagonist Bicuculline (0.5 mg/kg, i.p.) in D_1_-*CB_1_*-KO and WT littermates immediately after the NOR training phase. MK-801 did neither alter memory performance in WT mice nor did it rescue the amnesic phenotype of D_1_-*CB_1_*-KO littermates in the NOR task (**Figure 3A**; **S2A and S2B**). Conversely, Bicuculline completely reversed the memory impairment of D_1_-*CB_1_*-KO mice when injected immediately after training or 1h later, without affecting the NOR memory performance in WT littermates (**Figure 3A**; **S2A and S2B**).

**FIGURE 3.**
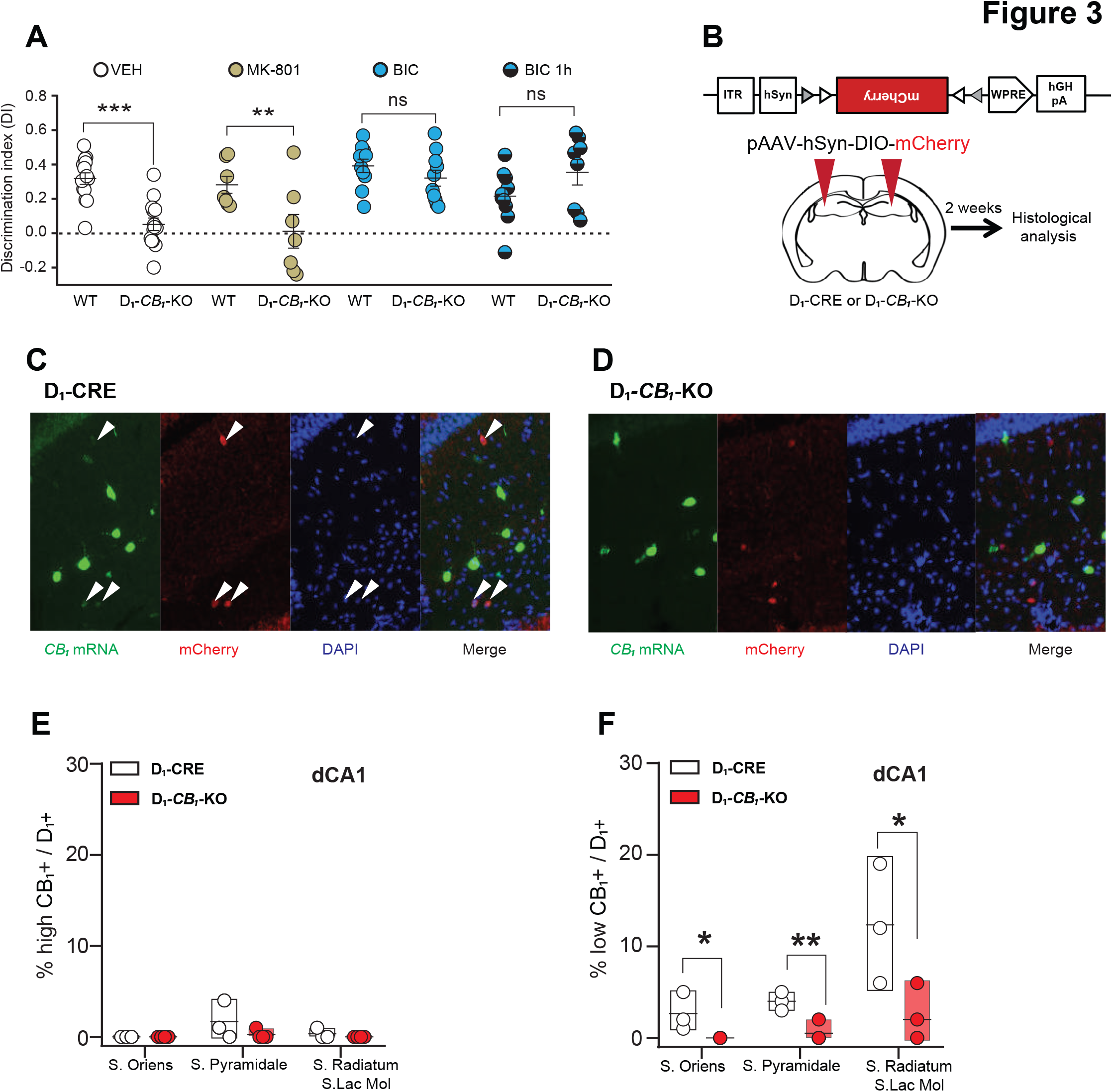
HIPPOCAMPAL CB_1_/D_1_-POSITIVE INTERNEURONS MODULATE SYNAPTIC GABAERGIC TRANSMISSION. (A) NOR memory performance of mutant mice administered with: vehicle [n(D_1_-*CB_1_*-WT)=14 and n(D_1_-*CB_1_*-KO)=14], MK-801 [0.1 mg per kg, intraperitoneal; n(D_1_-*CB_1_*-WT)=7 and n(D_1_-*CB_1_*-KO)=7], Bicuculline immediately after [n(D_1_-*CB_1_*-WT)=10 and n(D_1_-*CB_1_*-KO)=10] or 1 hour after the training phase [n(D_1_-*CB_1_*-WT)=10 and n(D_1_-*CB_1_*-KO)=8]. (B) Schematic representation of the experimental procedure to detect CB1 mRNA in D1-positive cells. (C and D) Representative images from the dorsal hippocampal CA1 region of D_1_-CRE and D_1_-*CB_1_*-KO expressing mCherry in D_1_-positive cells, after fluorescence *in situ* hybridization labelling of *CB_1_* mRNA (in green) and immunolabeling of mCherry protein (in red).(C) Representative pictures of D_1_-CRE mice expressing mCherry (in red) in D_1_-positive cells and *CB_1_* mRNA. White arrows indicate colocalization between CB_1_-positive and D_1_-positive cell bodies. (D) Representative pictures of D_1_-*CB_1_*-KO mice expressing mCherry (in red) in D_1_-positive cells and *CB_1_* mRNA. (E) Floating bars indicating the layer specific distribution of the % of cell bodies expressing high amounts of CB_1_ which colocalize with mCherry-positive (i.e. D_1_-positive) in D_1_-CRE and D_1_-*CB_1_*-KO. (F) Floating bars indicating the layer specific distribution of the % of cell bodies expressing low amounts of CB_1_ which colocalize with mCherry-positive (i.e. D_1_-positive) in D_1_-CRE and D_1_-*CB_1_*-KO. Data, mean ± SEM. *p < 0.05, **p < 0.01, ns=not significant. See also **Figure S2** and **Table S1**.

These data indicate that excessive GABA_A_ activity is involved in the phenotype of D_1_-*CB_1_*-KO mice. A large proportion of GABAergic hippocampal interneurons contain CB_1_ mRNA, which is expressed at different levels [high CB_1_- and low CB_1_-expressing cells, (Marsicano and Lutz, 1999)]. Conversely, D_1_ mRNA is expressed at very low levels in the hippocampus (http://mouse.brain-map.org/experiment/show/35, data not shown), which makes it difficult to accurately quantify its expression above background. Therefore, in order to pinpoint which CB_1_-positive interneurons in the hippocampus contain D_1_ receptors, we combined fluorescence *in situ* hybridization for CB_1_ mRNA in D_1_-Cre and D_1_-*CB_1_*-KO mice carrying viral Cre-dependent expression of mCherry (see methods and **Figure 3B**). As described (Marsicano and Lutz, 1999), detectable levels of CB_1_ receptor mRNA were present throughout the hippocampus both in pyramidal neurons and in GABAergic interneurons (**Figure S2C)**. The distribution of mCherry-tagged D_1_-positive neurons in the dorsal CA1 region of D_1_-CRE mice was similar to previously published data (Puighermanal et al., 2017, Gangarossa et al., 2012). Double staining revealed that virtually no high CB_1_-expressing interneurons in strata oriens, pyramidale, radiatum or lacunosum moleculare contain D_1_ receptors (**Figure 3C-F**, **S2C**). Conversely, D_1_ receptor is present in a small subpopulation of low CB_1_-expressing interneurons in the stratum oriens (2.7±0.7%), in the stratum pyramidale (4.0±0.4%) and in the strata radiatum/lacunosum moleculare (12.3±3.8%, **Figure 3C,F**). Importantly, this co-expression was virtually abolished in hippocampi of D_1_-*CB_1_*-KO mice (**Figure 3C,D,F**).

Altogether, these data indicate that CB_1_-dependent modulation of a small subpopulation of D_1_-positive GABAergic interneurons is required during NOR memory consolidation.

### SYNAPTIC MECHANISMS UNDERLYING NOR MEMORY CONSOLIDATION AND ASSOCIATED HIPPOCAMPAL PLASTICITY

The data collected so far indicate that reduction of GABAergic signaling prevents the memory deficits of D_1_-*CB_1_*-KO mice, which are associated with an impairment of learning-induced plasticity in the CA1 region of the hippocampus. Therefore, we tested whether inhibition of GABA_A_ receptors could rescue the lack of learning-dependent LTP enhancement observed in D_1_-*CB_1_*-KO mice. D_1_-*CB_1_*-KO and WT littermates trained in the NOR task received Bicuculline or vehicle before testing LTP induction in hippocampal circuits. In vehicle-treated animals, D_1_-*CB_1_*-KO mice showed no training-induced LTP enhancement as compared to WT littermates (**Figure 4A-C**). Strikingly, whereas Bicuculline treatment did not affect LTP induction and maintenance in WT animals, the same treatment rescued the training-induced LTP of D_1_-*CB_1_*-KO mice to similar levels as in WT littermates (**Figure 4A-C**).

**FIGURE 4.**
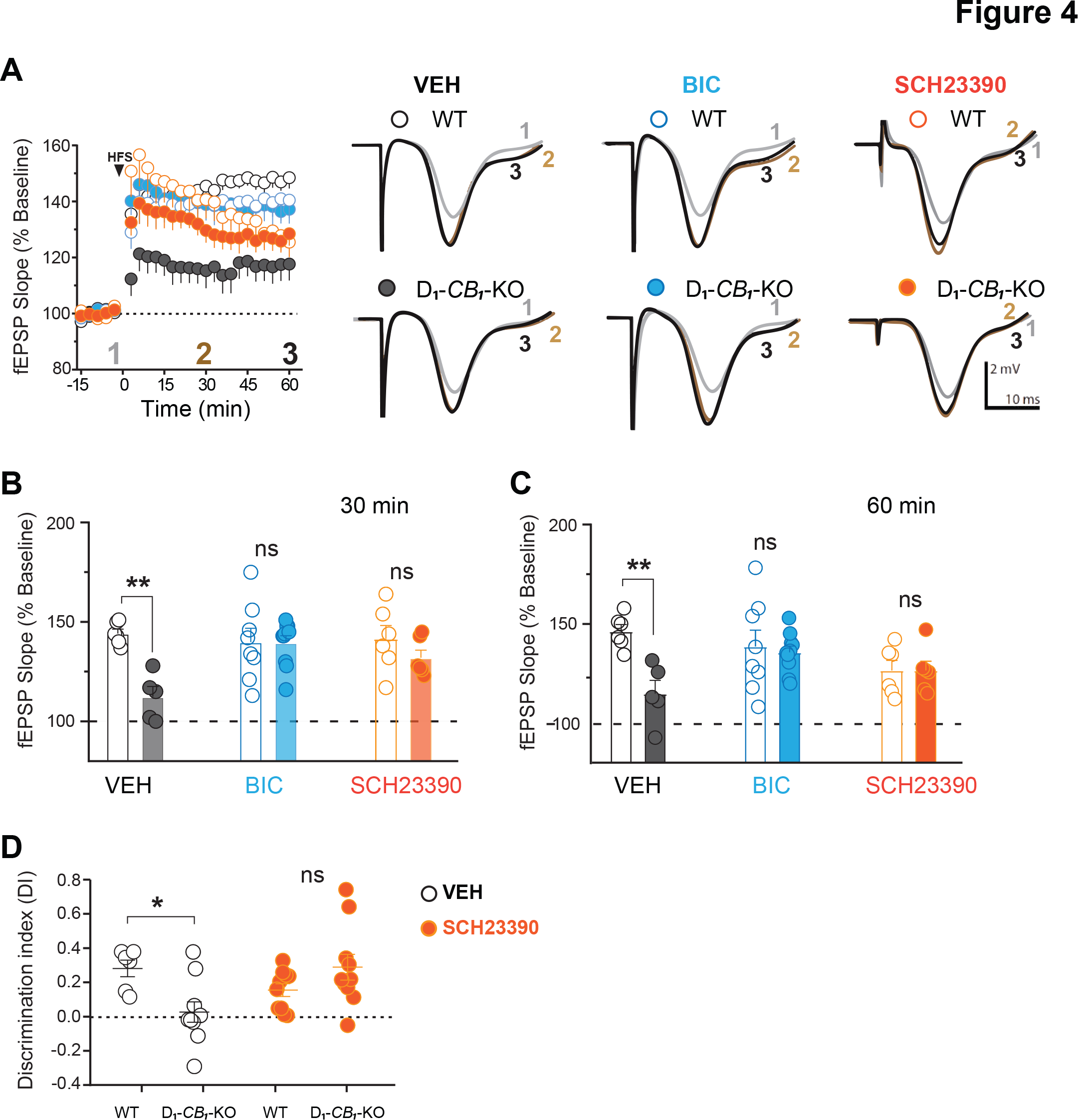
CELLULAR MECHANISMS LINKING D1-SIGNALLING WITH GABAERGIC ACTIVITY DURING LEARNING-INDUCED FACILITATION OF *IN VIVO* LTP AND MEMORY CONSOLIDATION. (A) GABA_A_ receptor antagonist Bicuculline, the D_1/5_ receptor antagonist SCH 23390 (0.3 mg per kg, intraperitoneal) on learning-induced modulation of *in vivo* LTP in D_1_-*CB_1_*-WT and D_1_-*CB_1_*-KO mice. Summary plots of recorded evoked fEPSP in Vehicle [n(D_1_-*CB_1_*-WT)=6 and n(D_1_-*CB_1_*-KO)=8], Bicuculline [0.5 mg per kg; intraperitoneal, n(D_1_-*CB_1_*-WT)=9 and n(D_1_-*CB_1_*-KO)=11] and SCH 23390[0.3 mg per kg; intraperitoneal, n(D_1_-*CB_1_*-WT)=6 and n(D_1_-*CB_1_*-KO)=6]. (B) Bar histograms of (A) representing normalized fEPSP from 30 minutes after HFS. (C) Bar histograms of (A) representing normalized fEPSP from 60 minutes after HFS. (D) Memory performance D_1_-*CB_1_*-WT and D_1_-*CB_1_*-KO mice after being injected with vehicle [n(D_1_-*CB_1_*-WT)=6 and n(D_1_-*CB_1_*-KO)=10] or SCH 23390 [0.3 mg per kg; intraperitoneal, n(D_1_-*CB_1_*-WT)=10 and n(D_1_-*CB_1_*-KO)=10] Traces on the right side of the summary plot (A) represent 150 superimposed evoked fEPSP before HFS (1, grey), 30 minutes (2, brown) and 60 minutes (3, black) after HFS. Data, mean ± SEM. *p < 0.05, **p < 0.01, ns=not significant. See also **Figure S3** and **Table S1**.

Recent data suggest that hippocampal D_1_-like receptors participate in memory formation, but little is known concerning the cell types involved in these processes (Lisman et al., 2011, Yamasaki and Takeuchi, 2017). To this point, our data indicate that CB_1_ receptor-dependent control of GABAergic transmission from a low number of hippocampal interneurons expressing D_1_ receptors is required to guarantee proper late consolidation of NOR memory. Therefore, it is possible that endocannabinoid actions are secondary to an activation of D_1_ receptors in these cells. To address this issue, we first reasoned that partial inhibition of D_1_ receptors should “replace” the lack of CB_1_-dependent control of neurotransmission in D_1_-*CB_1_*-KO mice. To test this idea, we administered a behaviorally sub-effective dose of the D_1/5_ antagonist SCH-23390 (SCH; 0.3 mg/kg; i.p.; **Figure S3A-S3C**) to D_1_-*CB_1_*-KO mice and WT littermates immediately after NOR acquisition and we analyzed the training-induced enhancement of *in vivo* LTP. This treatment slightly reduced the late phase of LTP in WT animals (**Figure 4A-C**). However, the antagonist abolished the differences between D_1_-*CB_1_*-KO mice and WT littermates (**Figure 4A-C**), indicating that reducing D_1_ receptor activity counteracts the absence of CB1 receptors in the mutants. If LTP is mechanistically linked to NOR consolidation, the same treatment should rescue the memory impairment of D_1_-*CB_1_*-KO mice. Vehicle-treated D_1_-*CB_1_*-KO animals displayed the previously observed memory impairment and the administration of SCH-23390 did not alter the behavior of WT mice (**Figure 4D and S4D, S4F**). Strikingly, however, the antagonist fully rescued the memory impairment of D_1_-*CB_1_*-KO littermates (**Figure 4D; S4D, S4F**).

Altogether, these results indicate that endocannabinoid-dependent regulation of hippocampal D_1_-positive interneurons represents a necessary step in the dopaminergic control of memory consolidation and associated synaptic plasticity.

## DISCUSSION

The present study reveals that a specific subpopulation of hippocampal CB_1_- and D_1_ receptor-positive GABAergic interneurons controls late consolidation of recognition memory and associated synaptic plasticity. By curbing inhibitory GABAergic activity at these interneurons, CB_1_ receptors participate in learning-induced facilitation of *in vivo* LTP and are required for consolidation of object recognition memory. Moreover, CB_1_ receptors in D_1_-positive neurons are necessary for physiological D_1_ receptor-dependent modulation of memory processes, indicating that these mechanisms are part of a complex regulatory circuit regulated by dopamine transmission in the hippocampus. By determining cellular and behavioral functions of a specific CB_1_-expressing interneuronal subpopulation, these data uncover an unforeseen role of CB_1_ receptors in the D_1_-dependent control of long-term memory.

Physiological or pharmacological modulation of the endocannabinoid system has a direct impact on episodic-like recognition memory processes *via* CB_1_ receptor-dependent control of different cell types within the hippocampus (Busquets-Garcia et al., 2017, Soria - Gomez et al., 2017). This multimodal influence of CB_1_ receptors on recognition memory raises the question about the differential cell type-, temporal- and spatial-specific contributions of the endocannabinoid system to memory processes. Past studies showed that ubiquitous (CB_1_-KO) or cell type-specific deletion of CB_1_ receptors from cortical glutamatergic (Glu-*CB_1_*-KO), forebrain GABAergic (GABA-*CB_1_*-KO), noradrenergic (DBH-*CB_1_*-KO) or serotonergic (TPH-*CB_1_*-KO) neurons did not impact NOR memory (Puighermanal et al., 2009, Busquets-Garcia et al., 2016), whereas their deletion in astrocytes does (Robin et al., 2018). Surprisingly, in the present study we observed that CB_1_ present in a specific subpopulation of hippocampal GABAergic interneurons expressing D_1_ receptors impairs the transition from short- to long-term memory processes. These apparently contrasting observations underline the complexity of cannabinoid control of learning and memory in the hippocampus.

The fact that the deletion of CB_1_ receptors in all body cells or in all forebrain GABAergic neurons does not reproduce the phenotype of D_1_-*CB_1_*-KO mice (Puighermanal et al., 2009) can be explained by different possibilities. Long-term deletion of the *CB_1_* gene starting from early developmental stages both in *CB_1_*-KO and GABA-*CB_1_*-KO mice might induce compensatory mechanisms (El-Brolosy and Stainier, 2017, El-Brolosy et al., 2019) masking the functional role of the receptor in NOR memory. An alternative or complementary explanation might point to the presence of different subpopulations of brain cells expressing CB_1_ and exerting opposite effects on memory processes. For instance, endocannabinoid signaling might promote or inhibit memory formation when acting at D_1_-positive cells or at other GABAergic subpopulations, respectively. Future studies will further dissect the functional impact of CB_1_ receptor signaling in different interneuronal subpopulations on NOR memory and other hippocampal functions.

Hippocampal-dependent memory functions are supported by interneuronal activity ensuring adequate spatiotemporal balance between excitation and inhibition of neuronal circuits (Klausberger and Somogyi, 2008). Amongst the known 21 different classes of GABAergic hippocampal interneurons, those expressing CCK contain the majority of CB_1_ receptors present in hippocampal cells (Harris et al., 2018, Marsicano and Lutz, 1999, Katona et al., 1999). Interestingly, while classically considered as a single class, it is currently known that CCK-positive interneurons encompass at least five cellular subtypes targeting different hippocampal *strata* and expressing specific markers (Harris et al., 2018). In our study, we found that D_1_ expression is present in a small sub-fraction of hippocampal interneurons containing CB_1_ mRNA. Very different levels of expression of the receptor characterize CB_1_-positive hippocampal interneurons, with a minority containing extremely high amounts and a majority much lower ones (Marsicano and Lutz, 1999). Interestingly, only a low percentage (5-15%) of low CB_1_-expressing interneurons contain D_1_ receptors, indicating that a very specific action of endocannabinoids is required to provide optimal consolidation of NOR memory. Therefore, our data reveal a specific function of a novel class of CB_1_- and D_1_-positive interneurons (Puighermanal et al., 2017).

We have previously shown that astroglial CB_1_ receptors are also necessary for consolidation of object recognition memory by allowing D-Serine availability at glutamatergic synapses (Robin et al., 2018). However, our data suggest that coordinated and temporally regulated actions of CB_1_ receptors at different cell types are required for NOR memory consolidation. In contrast to D_1_-*CB_1_*-KO animals that are selectively impaired in long-term NOR memory, mice lacking CB_1_ from astrocytes (GFAP-*CB_1_*-KO) lack both short- and long-term memory. In addition, D-serine rescues the lack of NOR memory in GFAP-*CB_1_*-KO mice only when administered immediately after learning, but not 1h later (Robin et al., 2018). Conversely, Bicuculline is effective in rescuing the phenotype of D_1_-*CB_1_*-KO mice also when administered 1h after learning. These observations suggest that endocannabinoid control of astrocytes is likely involved in the initial phases of the memory formation, whereas the CB_1_-dependent inhibition of GABAergic D_1_-positive interneurons would determine later phases of NOR consolidation. This idea is reinforced by the fact that GFAP-*CB_1_*-KO mice do not express *in vivo* LTP even in basal “home-cage” conditions (Robin et al., 2018), whereas D_1_-*CB_1_*-KO mice only lack the specific facilitation of LTP induced by learning. Altogether, these observations allow speculating that at least two distinct temporal windows exist in the CB_1_-dependent control of NOR. First, astroglial CB_1_ receptors are necessary for the plastic processes to initiate the memory. Later, the endocannabinoid-dependent regulation of D_1_-positive interneurons is required to maintain the memory trace for longer periods.

In this context, we cannot fully exclude that deletion of CB_1_ receptors in D_1_-positive cells does not involve also astrocytes that might express low levels of the dopamine receptor, as described in other brain regions (Nagatomo et al., 2017). However, no conclusive anatomical evidence so far has been presented for the expression of D_1_ receptors in hippocampal astrocytes [(Chai et al., 2017, Zhang et al., 2014), but see (Jennings et al., 2017) for D_1/5_ receptors pharmacological experiments] and D_1_-positive cells were shown to be mainly GABAergic interneurons in this brain region (Puighermanal et al., 2017, Gangarossa et al., 2012). Moreover, our data indicate that CB_1_ receptors impact memory formation by reducing inhibitory GABAergic transmission. Although astrocytes can release GABA (Yoon and Lee, 2014), the activation of astroglial CB_1_ receptors has been so far associated to increases of gliotransmitter release (Oliveira da Cruz et al., 2016). Therefore, in the light of the present knowledge, the classical presynaptic negative control of neurotransmitter release by CB_1_ receptors at D_1_-positive interneurons represents the most parsimonious explanation of the endocannabinoid impact on late consolidation of NOR memory.

LTP of the CA3-CA1 pathway shares similar molecular and cellular mechanisms with behavioral expression of episodic memory-like processes (Morris, 2013). However, whereas deletion of CB_1_ receptors from D_1_-positive cells impairs NOR memory, the same manipulation does not impair *in vivo* LTP of hippocampal synaptic transmission in naïve animals. In agreement with previous evidence in other experimental conditions (Li et al., 2003, Lemon and Manahan-Vaughan, 2006), WT mice exposed to the NOR learning task display a facilitation of *in vivo* LTP at CA3-CA1 synapses as compared to animals exposed to the same environment without any learning. Importantly, this facilitation is absent in D_1_-*CB_1_*-KO mice, indicating that the endocannabinoid control of D_1_-positive hippocampal interneurons is recruited only after learning. This facilitation might be due to a “real” stronger synaptic transmission after learning or to a decrease of baseline synaptic activity (Lisman, 2017), which might be occluded in D_1_-*CB_1_*-KO mice. The fact that partial blockade of GABA_A_ receptors in trained WT mice does not alter the LTP facilitation, suggests that this phenomenon is due to a genuine increase of LTP. However, our data indicate that reducing GABAergic transmission in D_1_ positive cells is required for this form of learning-induced synaptic plasticity. Moreover, these results reinforce the idea (Lisman et al., 2011) that, in order to reveal relevant mechanisms, investigations on synaptic plasticity associated to memory processes should include not only naïve animals, but also behaviorally challenged ones.

D_1_ receptor activity in the hippocampus can modulate long-term memory and synaptic plasticity (Lisman et al., 2011, Yamasaki and Takeuchi, 2017). However, the cell types involved in these D_1_ receptor-dependent functions remain mostly unknown. Our data suggest that D_1_/CB_1_-positive hippocampal interneurons are one of the targets of dopaminergic control of learning and memory processes. Indeed, partial inhibition of D_1_-like receptors is able to rescue the phenotype of D_1_-*CB_1_*-KO mice, suggesting that D_1_/CB_1_-positive interneurons belong to a larger circuit allowing the dopaminergic control of memory. For instance, recent data indicate that locus coeruleus projections to the hippocampus can release both noradrenaline and dopamine, with the latter playing key roles in the coding of novelty and memory (Yamasaki and Takeuchi, 2017). Interestingly, it was recently shown that parvalbumin (PV)-expressing interneurons require D_1_-like receptor activity for ongoing consolidation up to 14h after learning (Karunakaran et al., 2016). Partial pharmacological blockade of GABA_A_ receptors rescues the memory phenotype of D_1_-*CB_1_*-KO mice when injected immediately or 1h after learning. The fast pharmacodynamic clearance of the GABA_A_ antagonist Bicuculline in rodents (Wu et al., 2013), suggests that the counteracting effect of this drug should be disappeared few hours after treatment, placing the mutant mice in similar conditions as in controls. Therefore, it is tempting to speculate that cannabinoid control of D_1_/CB_1_/CCK-positive interneurons might participate in the consolidation of memory few hours after learning, whereas D_1_/PV-positive interneurons might be engaged at later time points. Future studies will address this intriguing hypothesis using the same memory protocols.

Altogether, these data reveal that functionally distinct cell types are present in the general population of hippocampal GABAergic interneurons expressing CB_1_ receptors. In particular, D_1_/CB_1_-positive interneurons provide specific behavioral and hippocampal synaptic mechanisms sustaining the fine-tuned regulation of memory processes. The close interaction of CB_1_ and D_1_ receptors in modulating recognition memory might provide novel therapeutic frameworks for the treatment of cognitive diseases characterized by alterations of both or either endocannabinoid and dopaminergic systems.

## ACKNOWLEDGMENTS

We thank Delphine Gonzales, Nathalie Aubailly and all the personnel of the Animal Facilities of the NeuroCentre Magendie for mouse care. We thank the Biochemistry Platform of Bordeaux NeuroCampus for help. We also thank all the members of Marsicano’s lab for useful discussions, Virginie Morales for invaluable support. This work was funded by: INSERM, European Research Council (Endofood, ERC–2010–StG–260515 and CannaPreg, ERC-2014-PoC-640923, MiCaBra, ERC-2017-AdG-786467), Fondation pour la Recherche Medicale (FRM, DRM20101220445 to G.M; DT20160435664 to J.F.O.d.C.), the Human Frontiers Science Program, Region Aquitaine, Agence Nationale de la Recherche (ANR, NeuroNutriSens ANR-13-BSV4-0006 and ORUPS ANR-16-CE37-0010-01) and BRAIN ANR-10-LABX-0043, to GM; NIH/NIDA (1R21DA037678-01), European Regional Development Fund, European Union’s Horizon 2020 Research and Innovation Programme (Grant Agreement 686009); French State/Agence Nationale de la Recherche/IdEx (ANR-10-IDEX-03-02) and Eu-Fp7 (FP7-PEOPLE-2013-IEF-623638), to A.B-G; FRM (ARF20140129235), to L.B; Ikerbasque (The Basque Foundation for Science), MINECO (Ministerio de Economía y Competitividad) PGC2018-093990-A-I00, to E.S-G.

## AUTHOR CONTRIBUTIONS

JFOdC performed and analyzed *in vivo* electrophysiology and behavioral experiments. AB-G and ES-G performed and analyzed behavioral experiments. LB and AC contributed to experiments using viral vectors. GL contributed to behavioral and *in vivo* electrophysiology experiments. MV and FJ-K performed immunohistochemistry experiments. ZZ, MV helped with the analysis of the data. JFOdC, AB-G, GM and ES-G wrote the manuscript. FD contributed to the writing. JFOdC, AB-G, GM and ES-G conceived and supervised the whole project and supervise the writing of the manuscript. All authors edited and approved the manuscript.

## METHODS

### EXPERIMENTAL MODEL AND SUBJECT DETAILS

#### ANIMALS

All experimental procedures were approved by the ethical committee of the French Ministry of Higher Education, Research and Innovation (authorization APAFIS#18111). Maximal efforts were made to reduce the suffering of the animals.

8 to 14 weeks-old naïve male D_1_-*CB_1_*-KO and WT littermates were used (Monory et al., 2007). 8–14 weeks-old male C57BL/6Rj mice purchased from Janvier (France). 8–12 weeks-old D_1_-CRE mice breed in the animal facilities of the U1215 we also used. Animals were housed collectively under standard conditions of temperature and humidity in a day/night cycle of 12/12 hours (light on at 7 am). Animals that underwent surgery were kept in individual cages after the procedures to avoid conflict with their littermates. Food and water was provided *ad libitum*. All the experiments were performed during the light phase. Behavioral experiments were performed from 9 am to 3 pm. Electrophysiology experiments were performed from 8 am to 7 pm.

### METHOD DETAILS

#### DRUG PREPARATION AND ADMINISTRATION

Bicuculline, MK-801 and SCH-23390 were purchased from Merck (formerly Sigma-Aldrich, France) and were dissolved to their final concentration in physiological saline (NaCl 0.9%). The exogenous DREADD ligand clozapine-N-oxide CNO (2 mg/kg) was purchased from Tocris Bioscience (Bristol, UK) and dissolved in saline after gently mixing with a vortex. All drugs were injected intraperitoneally (i.p.) in a volume of 10 ml/kg. Vehicle in all the conditions was composed of physiological saline (NaCl 0.9%)injections.Novel object-recognition memory task

#### NOVEL OBJECT RECOGNITION MEMORY

We used the novel object recognition memory task in an L-maze (NOR) (Busquets-Garcia et al., 2013, Busquets-Garcia et al., 2011, Puighermanal et al., 2009, Puighermanal et al., 2013, Robin et al., 2018, Hebert-Chatelain et al., 2016).

The task took place in a L-shaped maze made of dark grey polyvinyl chloride made by two identical perpendicular arms (35 cm and 30 cm long respectively for external and internal L walls, 4.5cm wide and 15 cm high walls) placed on a white background (Busquets-Garcia et al., 2011, Puighermanal et al., 2009). The task occurred in a room adjacent to the animal house with a light intensity fixed at 50 lux. The maze was overhung by a video camera allowing the detection and offline scoring of animal’s behavior. The task consisted in 3 sequential daily trials of 9 minutes each. During the habituation phase (day 1), mice were placed in the center of the maze and allowed to freely explore the arms in the absence of any objects. The training phase (day 2) consisted in placing the mice again in the corner of the maze in the presence of two identical objects positioned at the extremities of each arm and left to freely explore the maze and the objects. The testing phase occurred 24 hours later (day 3): one of the familiar objects was replaced by a novel object different in its shape, color and texture and mice were left to explore both objects. The position of the novel object and the associations of novel and familiar were randomized. All objects were previously tested to avoid biased preference. Memory performance was assessed by the discrimination index (DI). The DI was calculated as the difference between the time spent exploring the novel (TN) and the familiar object (TF) divided by the total exploration time (TN+TF): DI=[TN-TF]/[TN+TF]. Memory was also evaluated by directly comparing the exploration time of novel and familiar objects, respectively. Object exploration was defined as the orientation of the nose to the object at less than 2 cm. Experienced investigators evaluating the exploration were blind of treatment and/or genotype of the animals. Pharmacological treatments were immediately administered after the training phase.

#### *IN VIVO* ELECTROPHYSIOLOGY IN ANESTHETIZED MICE

Experiments were performed as described in (Robin et al., 2018). Mice were anesthetized in a box containing 5% Isoflurane (Virbac, France) before being placed in a stereotaxic frame (Model 900, Kopf instruments, CA, USA) in which 1.0% to 1.5% of Isoflurane was continuously supplied via an anesthetic mask during the whole duration of the experiment. The body temperature was maintained at ±36.5°C using a homoeothermic system (model 50-7087-F, Harvard Apparatus, MA, USA) and the state of anesthesia was assessed by mild tail pinch. Before surgery, 100 ml of the local anesthetic lurocaine (vetoquinol, France) was injected in the scalp region. Surgical procedure started with a longitudinal incision of 1.5 cm in length aimed to expose Bregma and Lambda. After ensuring the correct alignment of the head, two holes were drilled in the skull for electrode placement. Electrodes were: a glass recording electrode, inserted in the CA1 stratum radiatum, and a concentric stimulating bipolar electrode (Model CBARC50, FHC, ME, USA) placed in the CA3 region. Coordinates were as follows: CA1 stratum radiatum: A/P 1.5, M/L 1.0, DV 1.20; CA3: A/P 2.2, M/L 2.8, D/V 1.3 (20 insertion angle). The recording electrode (tip diameter = 1–2 mm, 2-4 MΩ) was filled with a 2% pontamine sky blue solution in 0.5M sodium acetate. At first the recording electrode was placed by hand until it reached the surface of the brain and then to the final depth using a hydraulic micropositioner (Model 2650, KOPF instruments, CA, USA). The stimulation electrode was placed in the correct area using a standard manipulator. Both electrodes were adjusted to find the area with maximum response. *In vivo* recordings of evoked field excitatory postsynaptic potentials (fEPSPs) were amplified 1000 times and filtered (low-pass at 1Hz and high-pass 3000Hz) by a DAGAN 2400A amplifier (DAGAN Corporation, MN, USA). fEPSPs were digitized and collected on-line using a laboratory interface and software (CED 1401, SPIKE 2; Cambridge Electronic Design, Cambridge, UK). Test pulses were generated through an Isolated Constant Current Stimulator (DS3, Digitimer, Hertfordshire, UK) triggered by the SPIKE 2 output sequencer via CED 1401 and collected every 2 s at a 10 kHz sampling frequency and then averaged every 180 s. Test pulse intensities were typically between 40-250 μA with a duration of 50 ms. Basal stimulation intensity was adjusted to 30%–50% of the current intensity that evoked a maximum field response. All responses were expressed as percent from the average responses recorded during the 15 min before high frequency stimulation (HFS). HFS was induced by applying 3 trains of 100 Hz (1 s each), separated by 20 s interval. fEPSP were then recorded for a period of 60 min. C57BL6/NRj mice underwent this *in vivo* electrophysiology procedure after the training phase of NOR task. Also, where specified, D_1_-*CB_1_*-KO and D_1_-*CB_1_*-WT received an injection of Bicuculine (0.5 mg *per* kg, intraperitoneal) or SCH 23390 (0.3 mg per kg, intraperitoneal) or vehicle immediately after undergoing training in NORT and before being subjected to the *in vivo* electrophysiology procedure. At the end the experiment, the position of the electrodes was marked (recording area: iontophoretic infusion of the recording solution during 180 s at 20mA; stimulation area: continuous current discharge over 20 s at +20mA) and histological verification was performed *ex vivo*.

#### SURGERY AND VIRAL ADMINISTRATION

Mice were anesthetized in a box containing 5% Isoflurane (Virbac, France) before being placed in a stereotaxic frame (Model 900, Kopf instruments, CA, USA) in which 1.0% to 1.5% of Isoflurane was continuously supplied via an anesthetic mask during the whole duration of the experiment. For viral intra-HPC AAV delivery, mice were submitted to stereotaxic surgery (as above) and AAV vectors were injected with the help of a microsyringe (0.25 ml Hamilton syringe with a 30-gauge beveled needle) attached to a pump (UMP3-1, World Precision Instruments, FL, USA). Where specified, D_1_-*CB_1_*-WT and D_1_-*CB_1_*-KO mice were injected directly into the hippocampus (HPC) or striatum (STR) (0.5 μl per injection site at a rate of 0.5 μl per min), with the following coordinates: HPC, AP −1.8; ML ±1; DV −2.0 and −1.5; Striatum: AP −1.34; ML ±2.8; DV −1.84. Following virus delivery, the syringe was left in place for 1 minute before being slowly withdrawn from the brain. CB_1_^*flox/flox*^ mice were injected with rAAV-CAG-DIO (empty control vector) or AAV-CAG-DIO-*CB_1_* to induce repression of the CB1 gene in hippocampal or striatal D_1_-positive cells. *CB_1_* coding sequence was cloned in rAAV-CAG-DIO vector using standard molecular cloning technology. The coding sequence was cloned inverted in orientation to allow CRE-dependent expression of CB_1_ receptors (Atasoy et al., 2008). In another experiment, and using the same procedure as described as above, D_1_-*CB_1_*-WT and D_1_-*CB_1_*-KO mice were injected intra hippocampally (AP −1.8; ML ±1; DV −2.0 and −1.5), with pAAV-hSyn-DIO-hM4D(Gi)-mCherry or pAAV-hSyn-DIO-mCherry (addgene, USA). For anatomical experiments and using the same procedure as above, D_1_-CRE and D_1_-*CB_1_*-KO were injected intra hippocampally with pAAV-hSyn-DIO-mCherry. In this specific experiment, expression was allowed to take place for 2 weeks. For the remaining experiments, animals were used around 4-5 weeks after local infusions. Mice were weighed daily and individuals that failed to regain the pre-surgery body weight were excluded from the following experiments.

#### IMMUNOHISTOCHEMISTRY ON FREE-FLOATING SECTIONS

Mice were anesthetized with pentobarbital (Exagon, Axience SAS, 400 mg/kg body weight), transcardially perfused with phosphate-buffered solution (PBS 0.1M, pH 7.4) before being fixed with 4% formaldehyde. Then brains were extracted, embedded with sucrose 30% for 3 days and finally frozen in 2-methylbutane (Sigma-Aldrich, France) at −80°C. Free-floating frozen coronal sections (40 μm) were collected with a cryostat and placed in PBS at room temperature (RT). Sections were permeabilized in a blocking solution of 10% donkey serum, 0.3% Triton X-100 in PBS for 1 hour at RT. Then, sections were incubated with a rabbit antibody against the C-myc epitope tag (1:1000, BioLegend) overnight at 4°C. After several washes with PBS, slices were incubated for 2 hours with a secondary anti-rabbit antibody conjugated to Alexa 488 (1:500, Fisher Scientific) and then washed in PBS at RT. Finally, sections were incubated with 4’,6-diamidino-2-phenylindole (DAPI 1:20000, Fisher Scientific) for 5 minutes before being washed, mounted and coverslipped. All the antibodies were diluted in blocking solution.

#### COMBINED FLUORESCENT *IN SITU* HYBRIDIZATION (FISH)/IMMUNOHISTOCHEMISTRY (IHC) ON FREE-FLOATING FROZEN SECTIONS

Mice were anesthetized with pentobarbital (Exagon, Axience SAS, 400 mg/kg body weight), transcardially perfused with PBS (0.1M, pH 7.4) before being fixed with 4% formaldehyde. Their brains were extracted, embedded with sucrose 30% for 3 days and finally frozen in 2-methylbutane (Sigma-Aldrich) at −80°C. Free-floating frozen coronal sections were collected with a cryostat (40 μm) and placed in a Rnase free PBS solution (PBS 0.1M, pH 7.4) at RT.

The endogenous peroxidases were inactivated by adding 3% H_2_O_2_ for 30 min to the free-floating sections. All endogenous biotin, biotin receptors, and avidin binding sites present in the tissue were blocked by using the Avidin/Biotin Blocking Kit (Vector Labs, USA). Then, the slices were incubated overnight at 4°C with a rabbit polyclonal antibody against DsRed (1:1000, Takara Bio USA) diluted in a blocking solution of 0.3% Triton X-100 in PBS-DEPC. The following day, after several washes, the sections were incubated with a secondary antibody anti-rabbit conjugated to a horseradish peroxidase (HRP) (1:500, Cell Signaling Technology) and later with a Biotin Tyramide (1:100, PerkinElmer) for 10 min at RT. After several washes, the slices were fixed with 4% of formaldehyde for 10 min and placed with 0.2M HCl for 20 min at RT. Then, an acetylation step (10 min in 0.1 M Triethanolamine, 0.25% Acetic Anhydride) was performed to reduce non-specific probe binding. Digoxigenin (DIG)-labeled riboprobe against mouse CB_1_ receptor (1:1000) was prepared as described (Marsicano and Lutz, 1999). After hybridization overnight at 60°C with the probes, the slices were washed with different stringency wash buffers at 65°C. Then, the sections were incubated with 3% of H_2_O_2_ for 30 min at RT and blocked 1 hour with a blocking buffer prepared according to the manufacturer’s protocol. Anti-DIG antibody conjugated to HRP (1:2000, Roche) were applied for 2 hours at RT. The signal of CB1 hybridization was revealed by a tyramide signal amplification (TSA) reaction using FITC-labeled tyramide (1:80 for 12 min, Perkin Elmer). After several washes the free-floating slices were incubated overnight with Streptavidin-Texas Red (1: 400, PerkinElmer). Finally, the slices were incubated in DAPI (1:20000; Fisher Scientific), washed, mounted, coverslipped and visualized with an epifluorescence Leica DM 6000 microscope.

### QUANTIFICATION AND STATISTICAL ANALYSIS

#### DATA COLLECTION

No statistical methods were used to pre-determine sample sizes, but they are similar to those reported in previous publications. All data collection and/or analysis were performed blind to the conditions of the experimenter except for the *in vivo* electrophysiological experiments. All mice were assigned randomly to the different experimental conditions.

#### FLUORESCENCE QUANTIFICATIONS

Cells expressing mRNAs were quantified in the different layers (stratum oriens, stratum pyramidale, stratum radiatum and stratum lacunosum moleculare) of the dorsal hippocampus. CB1 positive cells were classified according to the level of transcript visualized by the intensity of fluorescence (Marsicano and Lutz, 1999; Terral et al., 2019). ‘‘High-CB1’’ cells were considered to be round-shaped and intense staining covering the entire nucleus whereas ‘‘Low-CB1’’ cells were defined with discontinuous shape and lowest intensity of fluorescence allowing the discrimination of grains of staining.

#### STATISTICAL ANALYSES

Data were expressed as mean ± SEM or single data points and were analyzed with Prism 6.0 (Graphpad Software), using two-tails *t-*test (paired, unpaired) or one-way ANOVA (Dunnett’s), two-way ANOVA (sidak’s).

