## Supplementary Figures and table for "Specific hippocampal interneurons shape consolidation of recognition memory"

### Figure S1

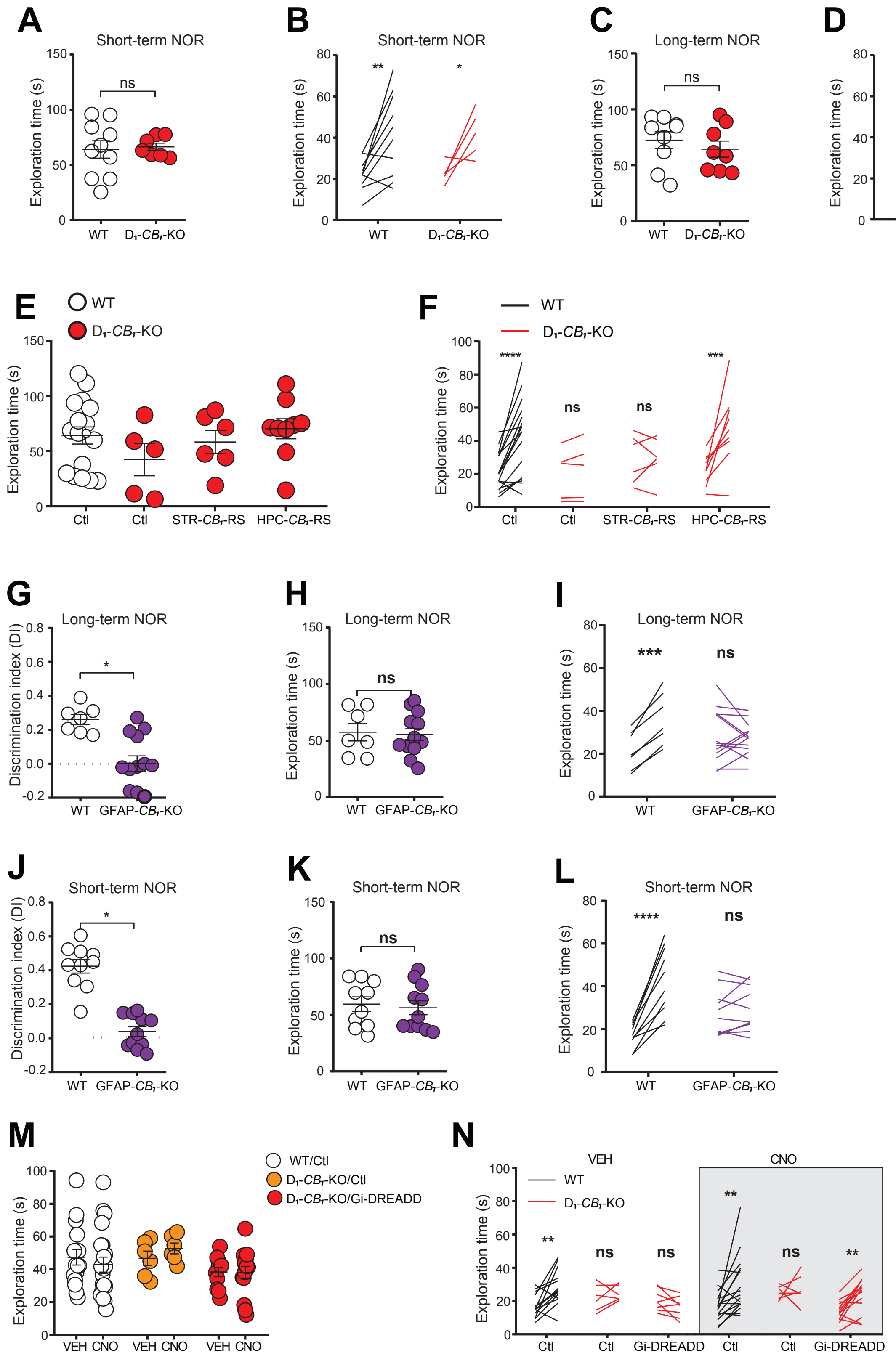

A
Short-term NOR
Exploration time (s)
ns
WT
D1-CB1-KO
B
Short-term NOR
Exploration time (s)
**
*
WT
D1-CB1-KO
C
Long-term NOR
Exploration time (s)
ns
WT
D1-CB1-KO
D
Long-term NOR
Exploration time (s)
****
ns
WT
D1-CB1-KO
E
WT
D1-CB1-KO
Exploration time (s)
Ctl
Ctl
STR-CB1-RS
HPC-CB1-RS
F
WT
D1-CB1-KO
Exploration time (s)
****
ns
ns
***
Ctl
Ctl
STR-CB1-RS
HPC-CB1-RS
G
Long-term NOR
Discrimination index (DI)
*
WT
GFAP-CB1-KO
H
Long-term NOR
Exploration time (s)
ns
WT
GFAP-CB1-KO
I
Long-term NOR
Exploration time (s)
***
ns
WT
GFAP-CB1-KO
J
Short-term NOR
Discrimination index (DI)
*
WT
GFAP-CB1-KO
K
Short-term NOR
Exploration time (s)
ns
WT
GFAP-CB1-KO
L
Short-term NOR
Exploration time (s)
****
ns
WT
GFAP-CB1-KO
M
WT/Ctl
D1-CB1-KO/Ctl
D1-CB1-KO/Gi-DREADD
Exploration time (s)
VEH CNO
VEH CNO
VEH CNO
N
VEH
CNO
WT
D1-CB1-KO
Exploration time (s)
**
ns
ns
**
ns
**
Ctl
Ctl
Gi-DREADD
Ctl
Ctl
Gi-DREADD

#### Figure S1. Related to Figure 1.

(A) Total exploration time, and (B) exploration time of the familiar versus the novel object of $D_1$-$CB_1$-KO mice and WT littermates in the Short-term NOR task. (C) Total exploration time, and (D) exploration time of the familiar versus the novel object of $D_1$-$CB_1$-KO mice and WT littermates in the Long-term NOR task. (E) Total exploration time, and (F) exploration time of the familiar versus the novel object of STR-$CB_1$-RS, HPC-$CB_1$-RS and control mice in the Long-term NOR task. (G) Memory performance, (H) total exploration, and (I) exploration time of the familiar versus the novel object of GFAP-$CB_1$-KO mice and WT littermates in the Long-term NOR task. (J) Memory performance, (K) total exploration, and (L) exploration time of the familiar versus the novel object of GFAP-$CB_1$-KO mice and WT littermates in the Short-term NOR task. (M) Total exploration time, and (N) exploration time of the familiar versus the novel object of $D_1$-$CB_1$-KO and WT littermates intra-hippocampally injected with hM4D(Gi) virus or mCherry, with VEH or CNO in the Long-term NOR task.

See also **Table S1**.

#### Figure S2

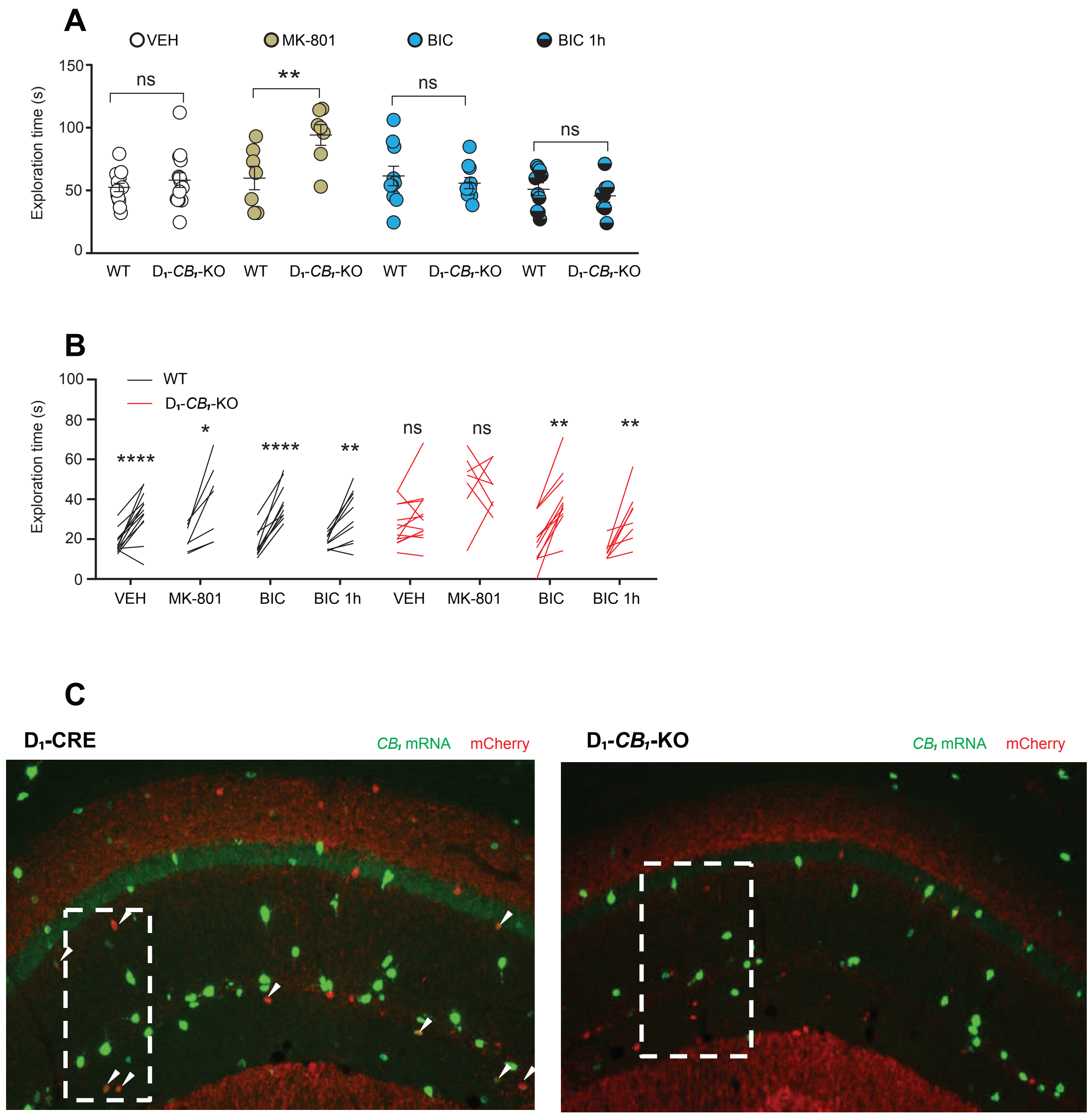
A
VEH
MK-801
BIC
BIC 1h
Exploration time (s)
150
100
50
0
ns
**
ns
ns
WT
D1-CB1-KO
WT
D1-CB1-KO
WT
D1-CB1-KO
WT
D1-CB1-KO
B
WT
D1-CB1-KO
Exploration time (s)
100
80
60
40
20
0
****
*
****
**
ns
ns
**
**
VEH
MK-801
BIC
BIC 1h
VEH
MK-801
BIC
BIC 1h
C
D1-CRE
CB1 mRNA
mCherry
D1-CB1-KO
CB1 mRNA
mCherry

#### Figure S2. Related to Figure 3

(A) Total exploration, and (B) exploration time of the familiar versus the novel object time of $D_1$-*$CB_1$*-KO mice and WT littermates with systemic injections of VEH, MK-801, and Bicuculine (BIC). (C) Representative micrographs of the dorsal hippocampus of $D_1$-CRE and $D_1$-*$CB_1$*-KO mice showing the co-expression of *$CB_1$* mRNA and mCherry ($D_1$ positive cells). The dotted white square is the area showed in main Figure 3C.

See also **Table S1**.

### Figure S3

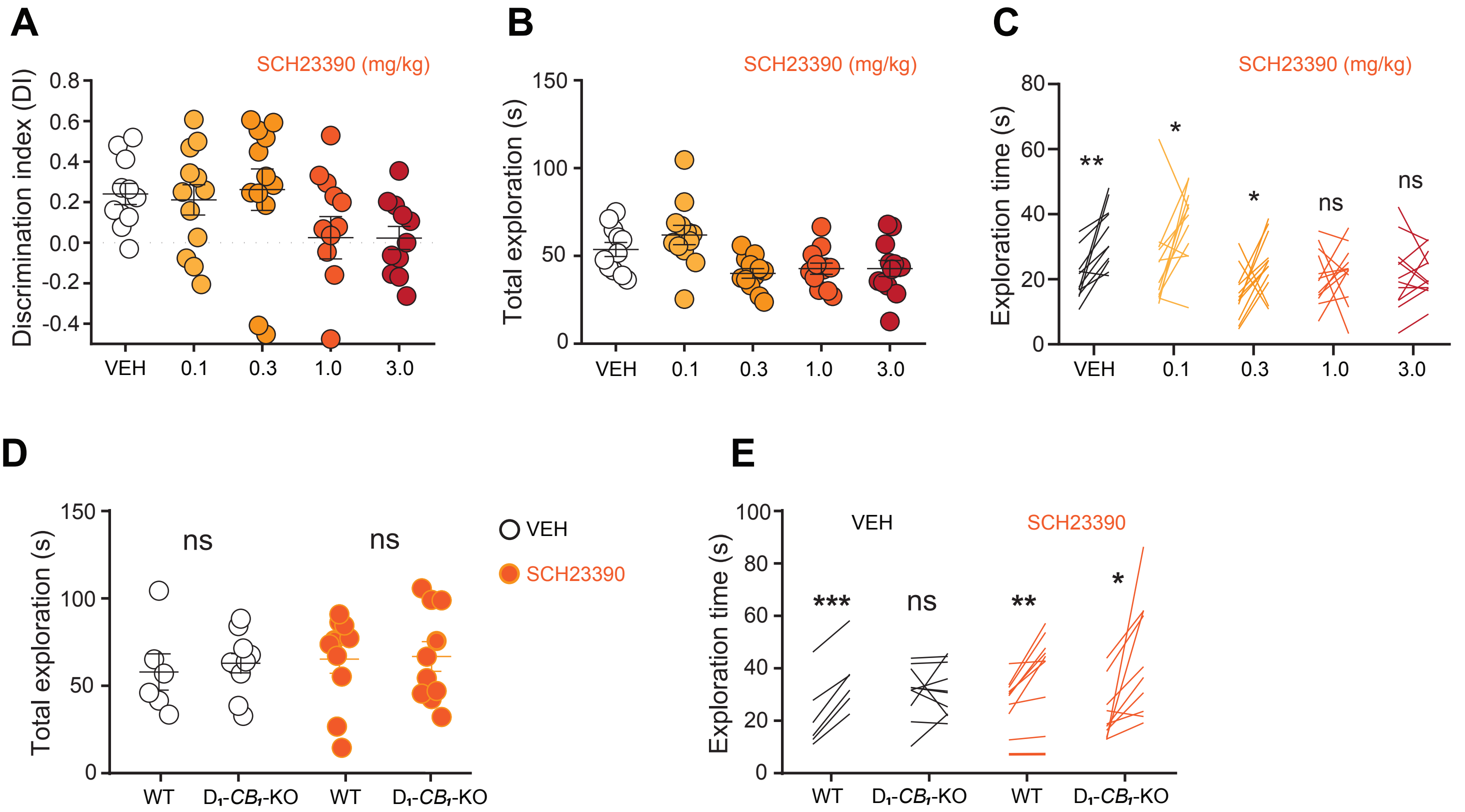
A
SCH23390 (mg/kg)
Discrimination index (DI)
0.8
0.6
0.4
0.2
0.0
-0.2
-0.4
VEH
0.1
0.3
1.0
3.0
B
SCH23390 (mg/kg)
Total exploration (s)
150
100
50
0
VEH
0.1
0.3
1.0
3.0
C
SCH23390 (mg/kg)
Exploration time (s)
80
60
40
20
0
**
*
*
ns
ns
VEH
0.1
0.3
1.0
3.0
D
Total exploration (s)
150
100
50
0
ns
ns
WT
D1-CB1-KO
WT
D1-CB1-KO
VEH
SCH23390
E
Exploration time (s)
100
80
60
40
20
0
VEH
SCH23390
***
ns
**
*
WT
D1-CB1-KO
WT
D1-CB1-KO

#### Figure S3. Related to Figure 4

(A) Memory performance, (B) Total exploration time, and (C) Exploration time of the familiar versus the novel object of C57BL6/NRj mice injected with different doses of SCH23390. (D) Total exploration, and (E) exploration time of the familiar versus the novel object time of $D_1$-$\mathit{CB_1}$-KO mice and WT littermates with systemic injections of SCH23390 (0.3 mg/kg).

See also **Table S1**.

### Table S1

| Figure | Panel | Conditions | "n" | Analysis (post-hoc test reported) | Factors analyzed | F-ratios | P values |
|---|---|---|---|---|---|---|---|
| 1 | B | D1-CB1-WT vs D1-CB1-KO | 10, 7 | Unpaired t-test | - | - | 0,708 |
| | C | D1-CB1-WT vs D1-CB1-KO | 9, 8 | Unpaired t-test | - | - | < 0,0001 |
| | F | D1-CB1-WT(Control) vs D1-CB1-KO (Control) | 17, 5 | Unpaired t-test | - | - | 0,0271 |
| | F | D1-CB1-KO: (Control) vs (R-CB1-STR) vs (R-CB1-HPC) | 5, 6, 9 | One-way ANOVA (Dunnet) | - | - | 0,0133 |
| | I | " VEH vs CNO": "Control", D1-CB1-KO-DREADD-Gi", " D1-CB1-KO-mCherry" | (16, 14) (11, 14) (6, 7) | 2-WAY ANOVA (Sidak) | Treatment vs Genotype | Interaction F (2, 62) = 5,107 | 0,0088 |
| | | | | | | Genotype F (2, 62) = 7,266 | 0,0015 |
| | | | | | | Treatment F (1, 62) = 4,587 | 0,0362 |
| 2 | B | D1-CB1-WT vs D1-CB1-KO - 30 min | 8, 8 | Unpaired t-test | - | - | 0,9444 |
| | | D1-CB1-WT vs D1-CB1-KO - 60 min | 8, 8 | Unpaired t-test | - | - | 0,4546 |
| | E | D1-CB1-WT vs D1-CB1-KO - 30 min | 8, 11 | Unpaired t-test | - | - | 0,0455 |
| | | D1-CB1-WT vs D1-CB1-KO - 60 min | 8, 11 | Unpaired t-test | - | - | 0,0253 |
| | G | D1-CB1-WT vs D1-CB1-KO - 30 min | 10, 10 | Unpaired t-test | - | - | 0,0266 |
| | | D1-CB1-WT vs D1-CB1-KO - 60 min | 10, 10 | Unpaired t-test | - | - | 0,0306 |
| 3 | A | " D1-CB1-WT vs D1-CB1-KO": "VEH" / "MK-801" / "BIC" / "BIC 1h" | (14, 14) | 2-WAY ANOVA (Sidak) | Treatment vs Genotype | Interaction F (3, 72) = 7,124 | 0,0003 |
| | | | (7, 7) | | | | |
| | | | (10, 10) | | | Treatment F (3, 72) = 6,787 | 0,0004 |
| | | | (10, 8) | | | Genotype F (1, 72) = 10,17 | 0,0021 |
| | F | S Oriens: D1-CRE vs D1-CB1-KO | (3, 4) | Unpaired t-test | - | - | 0,0453 |
| | | S Pyramidale: D1-CRE vs D1-CB1-KO | (3, 4) | Unpaired t-test | - | - | 0,0059 |
| | | S Radiatum / L Moleculare: D1-CRE vs D1-CB1-KO | (3, 4) | Unpaired t-test | - | - | 0,0337 |
| 4 | B | " D1-CB1-WT vs D1-CB1-KO": "VEH" / "Bicuculline" / "SCH 23390" at 30 min | (6, 5) | 2-WAY ANOVA (Sidak) | Genotype vs Treatment | Interaction F (2, 35) = 4,670 | 0,0159 |
| | | | (8, 10) | | | Genotype F (1, 35) = 10,30 | 0,0028 |
| | | | (6, 6) | | | Treatment F (2, 35) = 2,492 | 0,0973 |
| | C | " D1-CB1-WT vs D1-CB1-KO": "VEH" / "Bicuculline" / "SCH 23390" at 60 min | (6, 5) | 2-WAY ANOVA (Sidak) | Genotype vs Treatment | Interaction F (2, 35) = 4,232 | 0,0226 |
| | | | (8, 10) | | | Treatment F (1, 35) = 6,108 | 0,0185 |
| | | | (6, 6) | | | Genotype F (2, 35) = 2,014 | 0,1487 |
| | D | " D1-CB1-WT vs D1-CB1-KO": "VEH"/"SCH 23390" | (6, 10) | 2-WAY ANOVA (Sidak) | Genotype vs Treatment | Interaction F (1, 32) = 10,15 | 0,0032 |
| | | | (10, 10) | | | Treatment F (1, 32) = 0,9844 | 0,3285 |
| | | | | | | Genotype F (1, 32) = 1,236 | 0,2746 |

| Figure | Panel | Conditions | "n" | Analysis (post-hoc test reported) | Factors analyzed | F-ratios | P values |
|---|---|---|---|---|---|---|---|
| S1 | A | D1-CB1-WT vs D1-CB1-KO | 10, 7 | Unpaired t-test | - | - | 0,8161 |
| | B | D1-CB1-WT: Familiar vs Novel | 10 | Paired t-test | - | - | 0,0072 |
| | B | D1-CB1-KO: Familiar vs Novel | 5 | Paired t-test | - | - | 0,0421 |
| | C | D1-CB1-WT vs D1-CB1-KO | 9, 8 | Unaired t-test | - | - | 0,4572 |
| | D | D1-CB1-WT: Before vs After | 9 | Paired t-test | - | - | <0,0001 |
| | D | D1-CB1-KO: Before vs After | 8 | Paired t-test | - | - | 0,4821 |
| | E | D1-CB1-KO: (Control) vs (R-CB1-STR) vs (R-CB1-HPC) | 5, 6, 9 | One-way ANOVA (Dunnet) | - | - | 0,2335 |
| | F | D1-CB1-WT(Ctl): Familiar vs Novel | 17 | Paired t-test | - | - | <0,0001 |
| | F | D1-CB1-KO(Ctl): Familiar vs Novel | 5 | Paired t-test | - | - | 0,2137 |
| | F | D1-CB1-KO(STR-$CB_1$-RS): Familiar vs Novel | 6 | Paired t-test | - | - | 0,8794 |
| | F | D1-CB1-KO(HPC-$CB_1$-RS): Familiar vs Novel | 9 | Paired t-test | - | - | 0,0064 |
| | G | GFAP-CB1-WT vs GFAP-CB1-KO | 7, 13 | Unpaired t-test | - | - | 0,001 |
| | H | GFAP-CB1-WT vs GFAP-CB1-KO | 7, 13 | Unpaired t-test | - | - | 0,8118 |
| | I | GFAP-CB1-WT: Familiar vs Novel | 7 | Paired t-test | - | - | 0,0004 |
| | I | GFAP-CB1-KO: Familiar vs Novel | 13 | Paired t-test | - | - | 0,6032 |
| | J | GFAP-CB1-WT vs GFAP-CB1-KO | 10, 11 | Unpaired t-test | - | - | <0,0001 |
| | K | GFAP-CB1-WT vs GFAP-CB1-KO | 10, 11 | Unpaired t-test | - | - | 0,7143 |
| | L | GFAP-CB1-WT: Familiar vs Novel | 10 | Paired t-test | - | - | <0,0001 |
| | L | GFAP-CB1-KO: Familiar vs Novel | 11 | Paired t-test | - | - | 0,2716 |
| | M | D1-CB1-WT(Ctl): VEH vs CNO | 16, 21 | Unpaired t-test | - | - | 0,5043 |
| | M | D1-CB1-KO(Ctl): VEH vs CNO | 6,6 | Unpaired t-test | - | - | 0,2957 |
| | M | D1-CB1-KO(Gi-DREADD): VEH vs CNO | 11, 14 | Unpaired t-test | - | - | 0,9387 |
| | N | D1-CB1-WT(Ctl): Vehicle: Familiar vs Novel | 16 | Paired t-test | - | - | 0,0017 |
| | N | D1-CB1-KO(Ctl): Vehicle: Familiar vs Novel | 6 | Paired t-test | - | - | 0,461 |
| | N | D1-CB1-KO(Gi-DREADD): Vehicle: Familiar vs Novel | 11 | Paired t-test | - | - | 0,0133 |
| | N | D1-CB1-WT(Ctl): CNO: Familiar vs Novel | 21 | Paired t-test | - | - | 0,0057 |
| | N | D1-CB1-KO(Ctl): CNO: Familiar vs Novel | 6 | Paired t-test | - | - | 0,701 |
| | N | D1-CB1-KO(Gi-DREADD): CNO: Familiar vs Novel | 14 | Paired t-test | - | - | 0,001 |
| S2 | A | " D1-CB1-WT vs D1-CB1-KO": "VEH" / "Mk-801" / "Bicuculline"/ "Bicuculline" 1h | 14, 14 | 2-WAY ANOVA (Sidak) | Genotype vs Treatment | Interaction: F (3, 72) = 3,058 | P=0,0336 |
| | | | 7, 7 | | | Genotype: F (1, 72) = 5,457 | P=0,0223 |
| | | | 10, 10 | | | Treatment: F (3, 72) = 6,508 | P=0,0006 |
| | | | 10, 8 | | | - | - |
| | B | D1-CB1-WT: Vehicle: Familiar vs Novel | 15 | Paired t-test | - | - | <0,0001 |
| | B | D1-CB1-WT: Mk-801: Familiar vs Novel | 7 | Paired t-test | - | - | 0,0129 |
| | B | D1-CB1-WT: Bicuculline: Familiar vs Novel | 10 | Paired t-test | - | - | <0,0001 |
| | B | D1-CB1-WT: Bicuculline 1h: Familiar vs Novel | 10 | Paired t-test | - | - | 0,0033 |
| | B | D1-CB1-KO: Vehicle: Familiar vs Novel | 14 | Paired t-test | - | - | 0,1869 |
| | B | D1-CB1-KO: MK-801: Familiar vs Novel | 7 | Paired t-test | - | - | 0,8454 |
| | B | D1-CB1-KO: Bicuculline: Familiar vs Novel | 10 | Paired t-test | - | - | 0,0034 |
| | B | D1-CB1-KO: Bicuculline 1h: Familiar vs Novel | 8 | Paired t-test | - | - | 0,0081 |
| S3 | C | Vehicle | 11 | Paired t-test | - | - | 0,0018 |
| | C | SCH 23390 0.1 mg/kg | 12 | Paired t-test | - | - | 0,0259 |
| | C | SCH 23390 0.3 mg/kg | 12 | Paired t-test | - | - | 0,0295 |
| | C | SCH 23390 1.0 mg/kg | 12 | Paired t-test | - | - | 0,768 |
| | C | SCH 23390 3.0 mg/kg | 12 | Paired t-test | - | - | 0,8451 |
| | D | " D1-CB1-WT vs D1-CB1-KO": "VEH" / "SCH23390" | 6, 10 | 2-WAY ANOVA (Tukey) | Genotype vs Treatment | Interaction: F (1, 32) = 0,04417 | P=0,8349 |
| | | | 10, 10 | | | Genotype: F (1, 32) = 0,4676 | P=0,4990 |
| | | D1-CB1-WT: Vehicle: Familiar vs Novel | 6 | Paired t-test | - | Treatment: F (1, 32) = 0,1547 - | P=0,6967 0,0002 |
| | | D1-CB1-KO: Vehicle: Familiar vs Novel | 10 | Paired t-test | - | - | 0,7749 |
| | E | D1-CB1-WT: SCH23390: Familiar vs Novel | 10 | Paired t-test | - | - | 0,0038 |
| | E | D1-CB1-KO: SCH23390: Familiar vs Novel | 10 | Paired t-test | - | - | 0,0143 |
| | E | | | | | | |
| | E | | | | | | |
